# HIPK2 regulates homology-directed DNA repair and PARP inhibitor sensitivity

**DOI:** 10.64898/2026.05.28.728554

**Authors:** Patrick Weyerhäuser, Pierre-Olivier Frappart, Georg Nagel, Stephan Krieg, Teodora Nikolova, Wynand P. Roos, Kaveri Raja, Sahar Karbassi, Steffen Dickopf, Magdalena C. Liebl, Daniela Pfeiffer, Huong Becker, Yang He, Markus Christmann, Matthias Altmeyer, Thomas G. Hofmann

## Abstract

Repair of DNA double-strand breaks (DSBs) by homologous recombination (HR) counteracts genome instability and carcinogenesis. Cancer cells frequently show defects in HR which can be therapeutically exploited by hypersensitivity to poly(ADP-ribose) polymerase inhibitor (PARPi) treatment. Here we identify an unforeseen function of HIPK2 in HR repair and PARPi sensitivity. HIPK2 accumulates at DSBs and associates with DSB repair factors at DNA damage foci. DSB recruitment of HIPK2 requires checkpoint kinase ATM activity. DNA repair pathway analysis revealed that HIPK2 depletion specifically impairs HR. Mechanistically, we found that HIPK2 binds BRCA1 and phosphorylates BRCA1 at Ser1191, a site that regulates damage-induced BRCA1 protein stability. Consistently, HIPK2 depletion or pharmacological inhibition of HIPK2 results in reduced BRCA1 protein levels, and sensitizes BRCA1-proficient cancer cells to IR damage and PARPi treatment. In sum, our results identify a role for HIPK2 in HR through regulating BRCA1 protein levels, and propose HIPK2 inhibition as a novel strategy to sensitize BRCA1-proficient cancer cells to PARPi treatment.

## INTRODUCTION

The DNA damage response (DDR) plays a fundamental role in human physiology and pathophysiology, and activates DNA repair upon repairable damage or the cell death response upon irreparable damage^1, 2^. DSBs are a major source of genomic instability and activate the DNA damage checkpoint kinase ATM, which regulates the DNA damage response (DDR) through a complex signalling network promoting either DSB repair or, upon irreparable damage, induction of the cell death response^1–4^. DSBs are mainly repaired by two cellular pathways, homologous recombination (HR) and non-homologous end-joining (NHEJ). While HR provides a high-fidelity repair mechanism and requires the sister chromatid as template, NHEJ provides a fast and highly-efficient DSB repair system with reduced fidelity and increased mutational frequencies ^1, 3^. HR repair of DSBs is restricted to the S/G2 phase of the cell cycle and involves different steps including recruitment of DNA repair factors and DNA end resection by the coordinated action of CtIP-Mre11-Rad50-NBS1, Dna2 and Exo1 nucleases, which generate single-stranded DNA bound by RPA. RPA is subsequently replaced by the Rad51 recombinase, which forms a nucleofilament with the assistance of the repair factors BRCA1, PALB2 and BRCA2 ^3, 4^. Rad51 recombinase is essential for homology search, strand invasion and DNA polymerization. Successful HR requires numerous additional factors including the RecQ family of helicases, nucleases and resolvase complexes to complete the process and disassemble the protein complexes^5^. The complexity of HR repair is incompletely understood and likely relevant regulators and modulators of the HR pathway are still missing. Interestingly, cancer cells frequently harbour BRCA1 or BRCA2 mutations which inactivate the HR pathway, and which can be therapeutically used in the clinics by treatment with poly-ADP-ribose-polymerase inhibitors (PARPi)^6,7^.

HIPK2 is a member of the DYRK kinase family^8^, which is activated in response to DNA breaks and stimulates DNA damage-induced cell death ^9, 10^. HIPK2 functions as haplo-insufficient tumour suppressor in mice and man^11, 12^, and its dysregulation is linked to human pathologies including neurodegeneration, kidney fibrosis and cancer^11–15^. Undamaged cells and keep HIPK2 protein levels low through targeted degradation mediated by E3 ubiquitin ligases Siah1^16^ and WSB-1 ^17^. Upon DSB formation, such as triggered by ionizing radiation (IR) or Doxorubicin treatment, HIPK2 is activated and stabilized^16, 18–20^ by a mechanism involving dissociation of Siah1 through ATM-mediated phosphorylation^16^, HIPK2 autophosphorylation^21^ and subsequent changes in HIPK2 conformation catalysed by the cis/trans isomerase Pin1^21, 22^. Upon irreparable DSB damage, HIPK2 phosphorylates p53 at Ser46^10, 18, 23–27^ and SIRT1 Ser682^19^ phosphorylation which stimulates expression of pro-apoptotic p53 target genes in association with the tumour suppressor PML^19, 25^.

Although the function of HIPK2 in response to irreparable DNA damage is well-established, it remains currently unclear whether HIPK2 may also participate in DSB repair in less severe conditions. Here, we identify an unforeseen role of HIPK2 in the regulation of homology-directed DSB repair and PARP inhibitor sensitivity through interaction with BRCA1.

## RESULTS

### HIPK2 accumulates at DSBs

HIPK2 is a well-established regulator of p53-induced cell death in response to irreparable DNA damage ^10, 16, 18, 19, 21, 23, 28^, but its function in response to repairable DNA damage remains largely unclear. To investigate the role of HIPK2 in response to repairable DNA damage, we used colony formation assays to establish experimental conditions corresponding to repairable versus irreparable DNA damage induced by ionizing radiation (IR), a well-known inducer of DSBs (**Fig.1a**). Next, we used immunoblotting to follow HIPK2 protein stabilization and its activity on p53 phosphorylation using repairable versus irreparable IR damage. As expected^18, 29^, in response to repairable IR damage (2.5 Gy) HIPK2-mediated p53 Ser46 phosphorylation remained virtually absent (**Fig.1b**). In contrast, upon irreparable IR damage, p53 Ser46 phosphorylation as well as caspase-mediated PARP cleavage were clearly observed (**Fig.1b**), indicative for DNA damage-induced apoptosis activation. Interestingly, HIPK2 stabilization^16^ was detected in response to repairable DNA damage to a comparable extend as upon irreparable IR damage (**Fig.1b**), which is in line with HIPK2 stabilization through the ATM signaling pathway ^16, 18^ . Moreover, accumulation of HIPK2 in response to repairable IR damage pointed to a potential role of HIPK2 in DSB repair.

**Figure 1.**
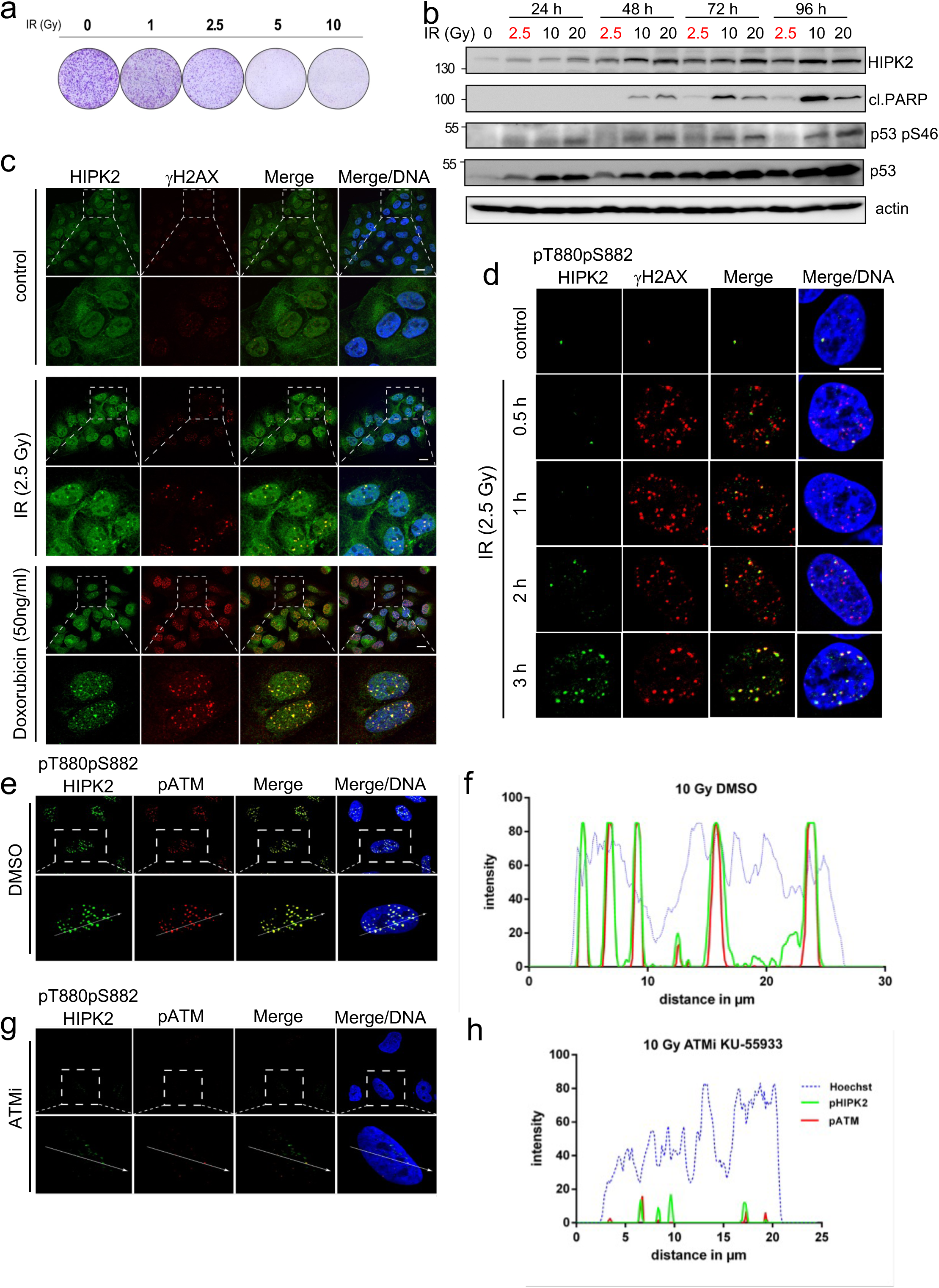
HIPK2 is activated upon repairable DSB damage and localizes to DSBs. **(a)** Colony formation analysis of U2OS cells irradiated with the indicated IR doses (Gy). A representative experiment is shown (n=3). **(b)** HIPK2 is stabilized upon repairable IR damage, in the absence of p53 Ser46 phosphorylation apoptosis. Immunoblot analysis of U2OS cells treated with a repairable IR dose (2.5 Gy, indicated in red colour) and irreparable IR damage (5, 10 and 20 Gy) as indicated. Cell lysates very analysed at different time points after IR treatment as indicated. Immunoblotting was performed with the antibodies indicated. A representative experiment is shown (n=2). **(c)** HIPK2 accumulates at DNA damage foci in response to IR and Doxorubicin treatment. U2OS cells were treated with IR (2.5 Gy) or Doxorubicin (50 ng/ml) or left untreated (control) as indicated. Cells were analysed by immunofluorescence staining using the antibodies indicated. DNA was stained with Hoechst (blue). Representative confocal images are shown (n=3). **(d)** Active HIPK2 phosphorylated at Thr880/Ser882 accumulates at DSBs upon IR damage. U2OS cells were treated with IR (2.5 Gy) or left untreated (control) as indicated. Cells were analysed by immunofluorescence staining using the antibodies indicated, and DNA was stained with Hoechst (blue). Representative confocal images are shown (n=3). **(e,g)** Active pHIPK2 colocalizes with active pATM at DSBs, and ATM activity is essential for recruitment of pHIPK2 to DSBs. U2OS cells were pre-treated with ATMi (g) for 1h or DMSO (e) and subsequently irradiated with 10 Gy IR. 4 hours later cells were fixed and analysed by immunofluorescence staining using the antibodies indicated. DNA was stained with Hoechst (blue). Representative confocal images are shown (n=3). **(f,h)** Intensity histogram analysis of the cells shown in (e,g,). Signal intensities of the indicated fluorescence channels (green: pHIPK2, red: pATM, blue: DNA) along the arrow indicated of DMSO-incubated or ATMi-treated U2OS cells after IR (10 Gy). Representative confocal images of a representative experiment (n=3) is shown.

To investigate whether HIPK2 may play a role in DSB repair, we first studied HIPK2 localization upon repairable IR damage, which triggers formation of DNA damage foci at DSBs. Strikingly, immunofluorescence staining and confocal microscopy indicated that HIPK2 partially colocalizes with DNA damage foci marked by γH2AX (**Fig.1c**). In addition, treatment with a none-cytotoxic amount of Doxorubicin, a topoisomerase II inhibitor leading to DSB formation, resulted in efficient recruitment of HIPK2 at DSBs (**Fig.1c**), indicating that HIPK2 is part of DNA damage foci, which play a role in DSB repair. Since HIPK2 stabilization and activation upon DNA damage requires site-specific auto-phosphorylation at Thr880/Ser882^21^, active HIPK2 can be visualized using phospho-specific pThr880/pSer882 HIPK2 antibodies ^21^. Indeed, confocal microscopy revealed that Thr880/pSer882-phosphorylated HIPK2 accumulates in a time-dependent manner at DSBs, where it colocalized with γH2AX (**Fig.1d**). These results indicate that active HIPK2 resides at DSBs. Furthermore, the activated DNA checkpoint kinase ATM phosphorylated at Ser1981, which is known to reside at DSBs^30^, colocalized with active HIPK2 at DSBs (**Fig.1e**). Intensity histogram analysis indicted that pHIPK2 and pATM show strong, but not complete colocalization (**Fig.1f**). Since ATM plays a central role for HIPK2 stabilization and activation in response to IR damage ^16, 18, 21^, we next addressed the role of ATM activity for DSB accumulation of HIPK2 by using an ATM inhibitor (ATMi). ATMi efficiently prevented the accumulation of HIPK2 and of activated pATM at DSBs (**Fig.1g,h**), illustrating a role of ATM for DSB recruitment of HIPK2. In sum, our results indicate that HIPK2 accumulates at DSBs in response to IR damage in an ATM-regulated fashion.

### HIPK2 regulates DSB repair by HR

Having demonstrated that HIPK2 accumulates at DSBs, we next determined its potential role in DSB repair by using a well-established nuclease-based reporter system for DSB repair through the NHEJ pathway (pimEJ5GFP)^31^ and HR (DR-GFP)^32^. Strikingly, HIPK2 depletion resulted in a robust reduction in HR efficiency (**Fig.2a**), whereas NHEJ was not diminished upon HIPK2 depletion (**Fig. 2b**). These results suggest that HIPK2 plays a role in the regulation of HR. HIPK2 depletion did not significantly alter cell cycle distribution of the depleted cells (**Fig.2c**), excluding the possibility that cell cycle-dependent effects may interfere with our analysis. To further support our finding that HIPK2 regulates HR, we analysed the impact of HIPK2 RNAi on the formation of sister-chromatid exchanges (SCEs), which are regulated by recombination events (**Fig.2d**). In line with a role for HIPK2 in regulating HR, MMS-induced SCE formation was significantly reduced in MMS-treated HIPK2-depleted cells (**Fig.2e**). In addition, HIPK2 depletion also slightly reduced the numbers of spontaneous SCEs in the untreated cells (**Fig.2e**). These results suggest a role in regulating HR.

**Figure 2.**
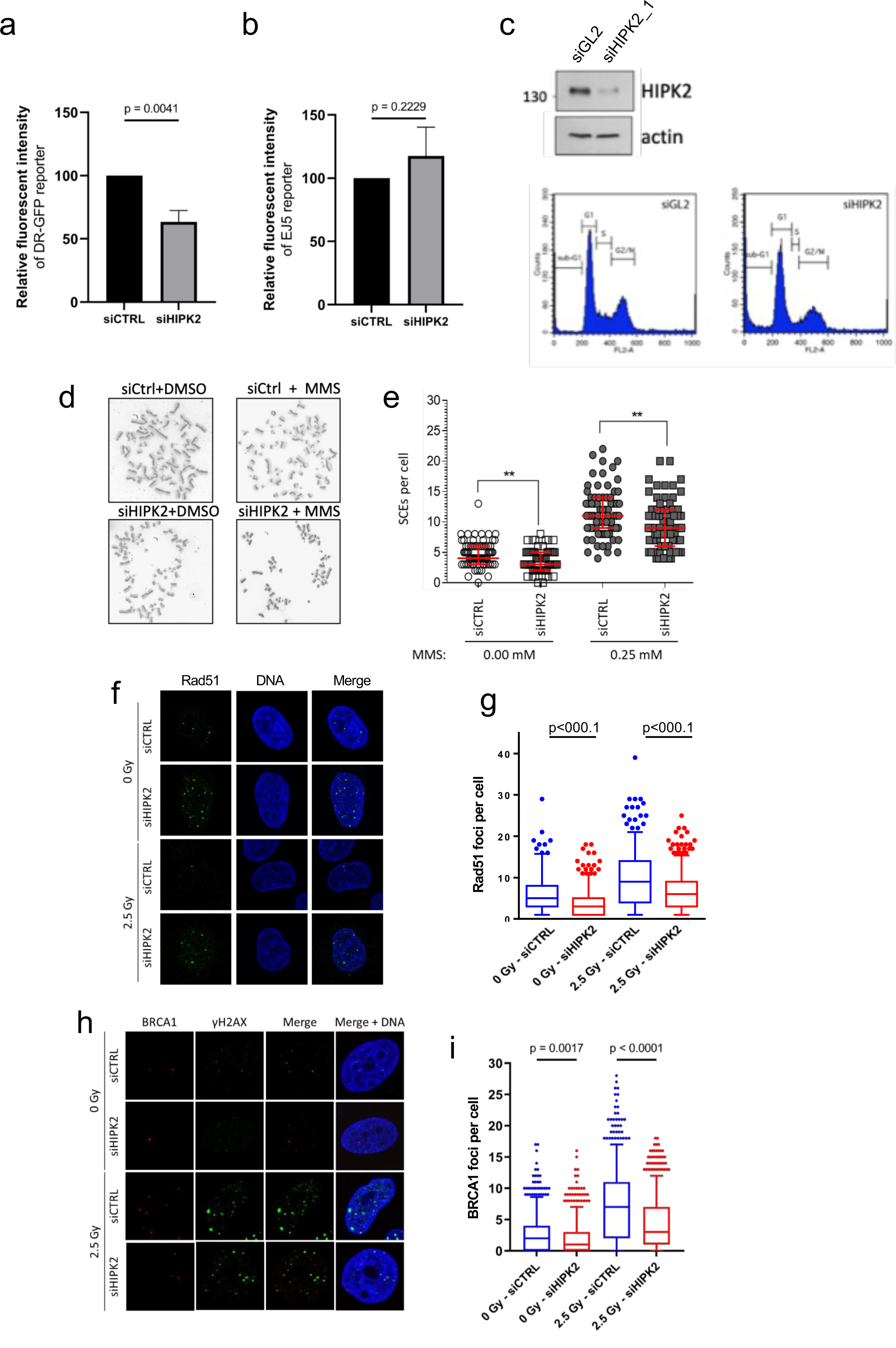
HIPK2 regulates homology-directed DSB repair. (a, **b)** HIPK2 is critical for HR repair, but not for NHEJ. U2OS cells stably expressing (a) the DR-GFP reporter plasmid or (b) the primEJ5-GFP reporter plasmid were depleted for HIPK2 as indicated, and 24 hours later transfected with the Sce-I endonuclease vector, and 24 hours later harvested and flow-cytometry analysis was performed. Statistical analysis was performed via paired and two-tailed t-test using the GraphPad Prism software. N=4 biological replicates. (a) Quantification and statistical analysis of DR-GFP reporter assays. (b) Quantification and statistical analysis of primEJ5-GFP reporter assays. **(c)** Immunoblotting demonstrating HIPK2 depletion in the DR-GFP reporter U2OS cells. A representative blot is shown (n=3). Lower panel: Flow-cytometry analysis showing cell cycle profile analysis of the cells depleted for HIPK2 in (c). **(d, e)** HIPK2 depletion results in reduced SCEs. (d) Metaphase spread of representative Sister-chromatid exchange (SCE) assays performed in control cells and Hipk2 depleted cells in the presence and absence of 0.25 µM MMS. (e) Quantification and statistical analysis of SCEs has been performed as described in the Methods section. (n=3 replicates, with at least 30 metaphases analysed each). **(f,g)** HIPK2 depletion leads to reduced Rad51 foci formation upon IR damage. Representative confocal images of Rad51 stainings in untreated and IR-treated siControl and siHIPK2 transfected U2OS cells are shown. (g) Quantification and statistical analysis Rad51 foci. More than 300 cells per condition were counted in total. More that 60 cells per condition were used for analysis. Box plots indicate 5-95 percentile, all other data points are shown as single dots. Statistical analysis was performed by unpaired and two-tailed t-test using GraphPad Prism. **(h,i)** HIPK2 depletion leads to reduced BRCA1 foci formation upon IR damage. (h) Representative confocal images of BRCA1 and γH2AX stainings in untreated and IR-treated siControl and siHIPK2 transfected U2OS cells are shown. (i) Quantification and statistical analysis BRCA1 foci. More than 300 cells per condition were counted in total. More that 60 cells per condition were used for analysis. Box plots indicate 5-95 percentile, all other data points are shown as single dots. Statistical analysis was performed by unpaired and two-tailed t-test using GraphPad Prism.

Foci formation of the Rad51 recombinase is a well-established read-out for the activation of the HR repair pathway. To further challenge our hypothesis on the role of HIPK2 in regulating HR, we next determined the effect of HIPK2 depletion on Rad51 foci numbers. HIPK2 depletion resulted in a significant reduction in both spontaneous as well as IR-induced Rad51 foci numbers (**Fig.2f,g**), further strengthening our hypothesis. Finally, we analysed the impact of HIPK2 depletion on formation of BRCA1 foci, which acts upstream of Rad51 foci formation in the HR pathway. HIPK2 knock-down resulted in a strong and significant reduction in BRCA1 foci numbers upon IR damage (**Fig.2h,i**), which supports our conclusion that HIPK2 plays a role in DSB repair through HR. In addition, our results suggest that HIPK2 might regulate HR through interplay with BRCA1.

### BRCA1 interacts with HIPK2

To test our hypothesis that HIPK2 may regulate HR through interaction with BRCA1, we analysed potential protein complex formation of HIPK2 and BRCA1 using immunoprecipitation analysis. The well-established HIPK2 binding partner p53 was used as a control^18, 23, 24^. In line with previous findings, p53 showed inducible complex formation with HIPK2 in response the irreparable DSB damage (0.75 µg|ml Doxorubicin), while almost no interaction was evident in undamaged cells (**Fig.3a**). Also, in cells treated with repairable DSB damage (50 ng/ml Doxorubicin) almost no p53 interaction was detectable (**Fig.3a**). Interestingly, BRCA1 already formed a complex with HIPK2 in undamaged cells, and complex formation was increased in response to repairable DSB damage (**Fig.3a**). In response to irreparable Doxorubicin treatment, BRCA1 complex formation with HIPK2 was weakened (**Fig.3a**). Of note, as a negative control the unrelated protein kinase AKT was used. As expected, AKT did not form a complex with HIPK2 (**Fig.3a**). Our results suggest that BRCA1 and HIPK2 form a specific protein complex in undamaged cells and upon repairable DSB damage.

**Figure 3.**
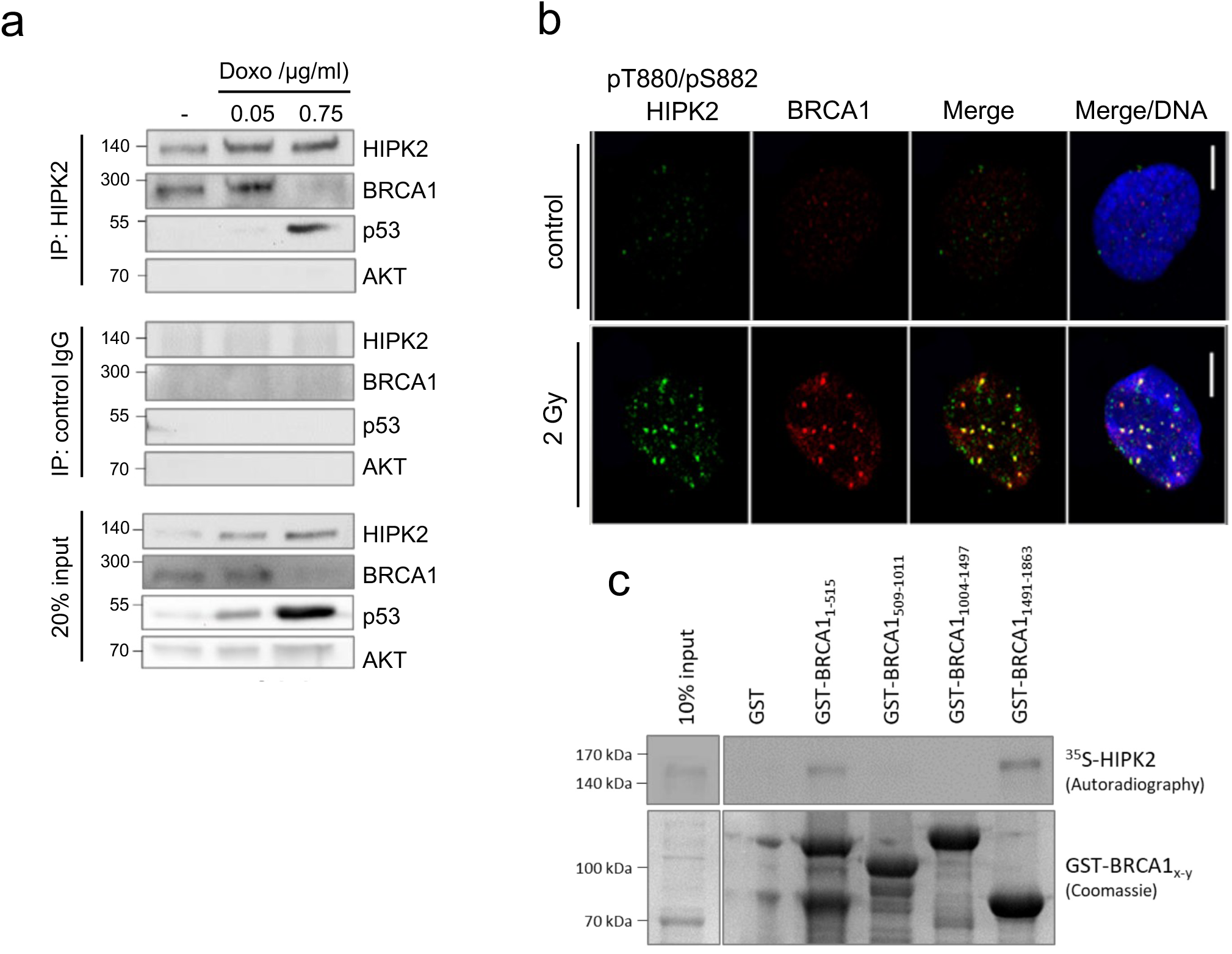
HIPK2 interacts with BRCA1. **(a)** BRCA1 forms a proteinncomplex with HIPK2 in cells. HCT116 cells treated with the indicated doses of Doxorubicin, or left untreated, were lysed after 18 hours and HIPK2 antibody or control IgG antibody IP was performed as indicated. Immunocomplexes were analysed by immunoblotting using the antibodies indicated. A representative experiment is shown (n=2 replicates). **(b)** BRCA1 colocalizes upon IR damage with active, pThr880/pSer882-phosphorylated HIPK2 at DSBs. Immunofluorescence staining was performed using the antibodies indicated, and representative confocal images are shown. DNA is stained with Hoechst (blue). N=3 replicates. **(c)** *In vitro* binding of BRCA1 and HIPK2. GST pulldown assays were performed using the indicated bacterially expressed GST-BRCA1 protein fragments spanning the entire BRCA1 protein along with full-length *in vitro* translated, ^35^S-labelled HIPK2 protein. GST-protein complexes were pulled down, separated by SDS-PAGE and analysed by autoradiography (upper panel). Loading was determined by Coomassie staining (lower panel). A representative experiment is shown (n=2).

Next, we determined BRCA1 and HIPK2 localization in response to DSB damage using confocal microscopy. Strikingly, in response to IR damage BRCA1 readily colocalized with HIPK2 at DSB sites (**Fig.3b**), while no clear colocalization was evident in undamaged cells. These results support an interaction of BRCA1 and HIPK2 upon DSB damage.

To investigated whether BRCA1 may physically bind to HIPK2, we performed GST pulldown assays using various bacterially expressed GST-BRCA1 fusions spanning the entire BRCA1 protein along with *in vitro* translated, ^35^S-labelled HIPK2. Strikingly, HIPK2 was pulled down with GST-BRCA1 aa1-515 and GST-BRCA1 aa1491-1863, suggesting two independent *in vitro* HIPK2-binding sites on BRCA1 (**Fig.3c**). Our results indicate that BRCA1 and HIPK2 colocalize upon repairable DSB induction and form a protein complex, and our data indicate that HIPK2 can physically interact with BRCA1.

### HIPK2 phosphorylates BRCA1 at Ser1191

HIPK2 is a Ser/Thr protein kinase mediating site-specific phosphorylation of numerous of its binding partners ^16, 19, 23, 26, 33, 34^. Therefore, we investigated whether BRCA1 is a HIPK2 substrate protein. To this end, we performed in *vitro kinase* assays in presence of ^32^P-labelled γATP using recombinant, bacterially expressed HIPK2 and recombinant, bacterially expressed GST-BRCA1 protein fragments altogether spanning the entire BRCA1 protein. Autoradiography of the dried SDS-PAGE indicated HIPK2-mediated phosphorylation of BRCA1 fragment spanning amino acids (aa) 1004-1497 and aa1491-1863 (**Fig.4a**). Next, we aimed at mapping the detailed phosphorylation sites within these BRCA1 fragments by generating smaller GST-BRCA1 fragments of the respective stretches, which identified Ser1191 of BRCA1 as a major HIPK2 *in vitro* phosphorylation site (**Fig.4b**). In this current study we decided to focus on the identified BRCA1 Ser1191 phosphorylation site. The *in vitro* phosphorylation site(s) detected in the BRCA1 fragment 1491-1863 (**Fig.4a**) will be addressed independently in future.

**Figure 4.**
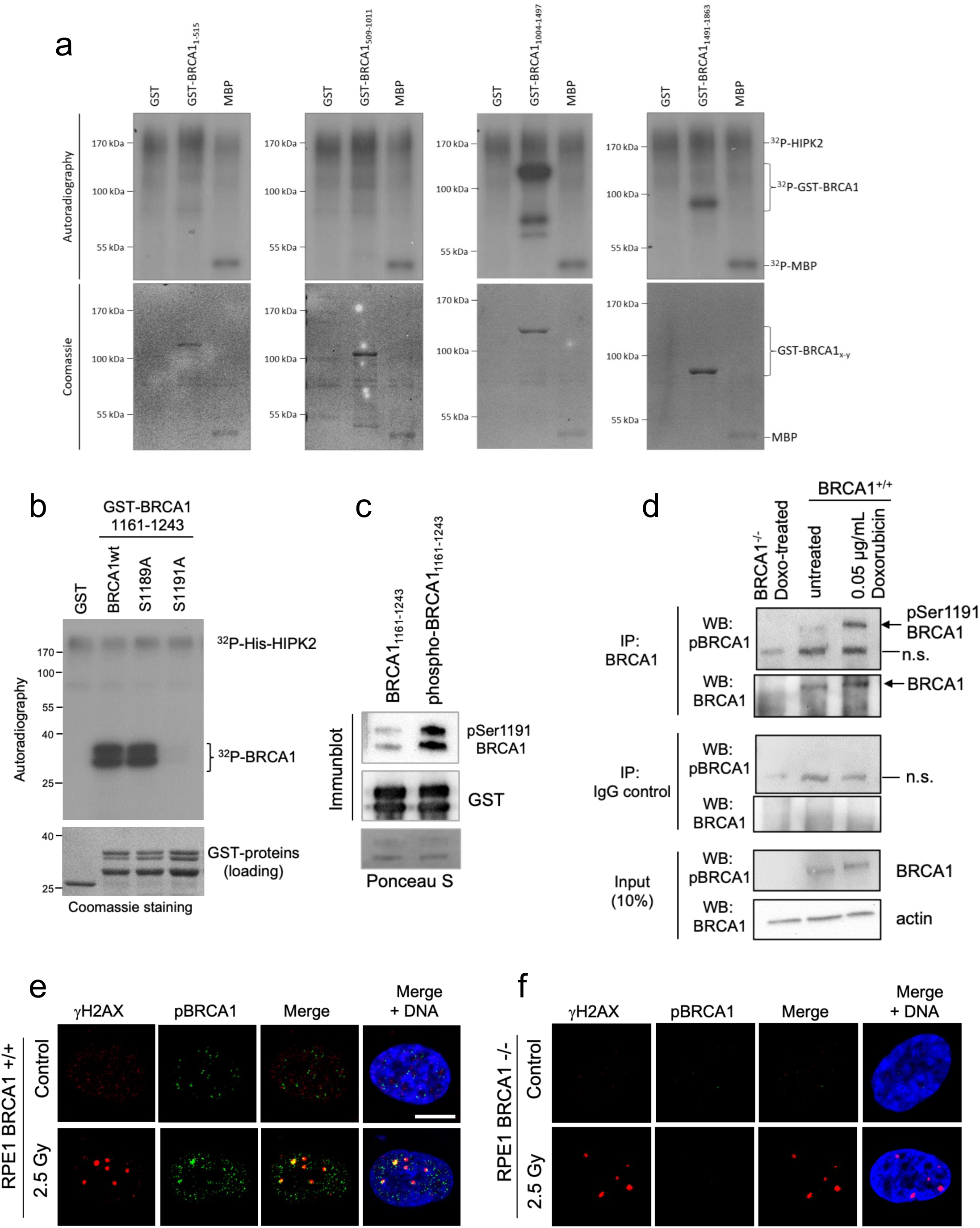
HIPK2 phosphorylates BRCA at Serine 1191. **(a)** HIPK2 phosphorylates BRCA1 *in vitro*. *In vitro* HIPK2-mediated phosphorylation of GST-BRCA1 protein fragment spanning the complete BRCA1 protein has been performed. *In vitro* kinase assays using bacterially expressed GST or the indicated GST-BRCA1 protein fragments along with bacterially expressed 6xHis-HIPK2 were performed, separated by SDS-PAGE, and dried gels were analysed by autoradiography (upper panel). MBP was used as a positive control as previously described^16, 23^. GST protein loading was determined by Coomassie staining (lower panel). A representative experiment is shown. **(b)** HIPK2 phosphorylates BRCA1 at Ser1191 *in vitro*. In vitro kinase assays were performed as described in (a) using the indicated GST-BRCA1 protein harbouring the indicated, site-specific mutations at potential phosphorylation sites. A representative experiment is shown. **(c)** An affinity-purified phosphorylation specific antibody raised against BRCA1 Ser1191 preferentially recognizes BRCA1 phosphorylated *in vitro* by HIPK2. Immunoblotting with affinity-purified BRCA1 pSer1191 antibodies using bacterially expressed GST and the indicated HIPK2-phosphorylated GST-BRCA1 fragment versus the unphosphorylated counterpart was performed. Protein loading and phosphorylation was validated using immunoblotting with indicated antibodies. **(d)** BRCA1 is phosphorylated at Ser1191 in response to DSB damage. BRCA1-deficient (BRCA1 -/-, p53 -/-) and BRCA1-proficient (BRCA1 +/+, p53 -/-) RPE-1 cells were treated with 0.05 µg/ml Doxorubicin and BRCA1 was immunoprecipitated as indicated. Immunoblotting was performed using the indicated antibodies. A representative experiment is shown. **(e, f)** Ser1191-phosphorylated BRCA1 accumulates at DSBs in response to IR damage in BRCA1 +/+ cells, but not in BRCA1 -/- cells. Immunofluorescence analysis has been performed using BRCA1 pSer1191 antibodies and γH2AX antibodies in e) BRCA1 +/+ RPE-1 cells and in f) BRCA1 -/- RPE-1 cells. DNA has been stained with Hoechst (blue). Representative images are shown.

Next, we aimed at investigating the HIPK2-catalysed BRCA1 pSer1191 phosphorylation in cells. Since no BRCA1 pSer1191 antibody against this site has been reported so far, we decided to raise a polyclonal phospho-specific antibody against the BRCA1 pSer1191 site. The antisera were affinity purified against the phospho-Ser1191 containing peptide and in the following against the corresponding none-phospho peptide to enrich for BRCA1 pSer1191-specific antibodies. Next, we validated the obtained antibody for its phospho-specificity. To this end, we *in vitro* phosphorylated a BRCA1 fragment using recombinant HIPK2 or left the fragment untreated, and determined the BRCA1 pSer1191 antibody reactivity to the phosphorylated and none-phosphorylated BRCA1 fragment using immunoblotting. Our results indicated that a strong preference of our pSer1191 BRCA1 antibody for recognizing the HIPK2-phosphorylated BRCA1 (**Fig.4c**), indicating phospho-specificity of the generated antibody.

Next, we used this novel tool to study BRCA1 phosphorylation at Ser1191 at the cellular level. To this end, cells were treated with a repairable dose of Doxorubicin, subsequently endogenous BRCA1 was immunoprecipitated from cell lysates and immunoblot analysis was performed. As a control, we used Doxorubicin-treated BRCA1 -/- cells. To this end, we used BRCA1-proficient (BRCA1 +/+) and BRCA1-deficient (BRCA1 -/-) RPE-1 cells^35^. BRCA1 Ser1191 phosphorylation was readily detected in Doxorubicin-treated BRCA1-proficient cells (**Fig.4d**). In addition, a weak pSer1191 BRCA1 signal was evident in untreated cells, which may suggest that Ser1191 does also occur at basal conditions and potentially contribute to BRCA1 stability in untreated cells. As expected, no pSer1191 BRCA1 signal was detected in BRCA1-deficient cells (**Fig.4d**), indicating BRCA1 specificity of the BRCA1 pSer1191 antibody

To investigate the subcellular localization of BRCA1 pSer1191, we performed immunofluorescence analysis. In BRCA1-proficient cells, Ser1191-phosphorylated BRCA1 was readily detected at DNA damage foci in response to IR damage, as indicated by partial colocalization with γH2AX (**Fig.4e**). In contrast, and as expected, IR-damaged BRCA1-deficient cells showed no signal for Ser1191 phosphorylated BRCA1, while γH2AX foci were readily detected in these cells (**Fig.4f**). In sum, these results indicate that HIPK2 phosphorylates BRCA1 at Ser1191 *in vitro* and in cells, and that BRCA1 pSer1191 is accumulating at DNA damage foci upon DSB induction.

### HIPK2 regulates BRCA1 protein levels

Phosphorylation of BRCA1 at Ser1191 has been recently reported to regulate BRCA1 stability upon IR damage through altering BRCA1 protein conformation by recruitment of the phospho-specific *cis/trans* isomerase Pin1^36^. These findings stimulated us to determine the effect of HIPK2 depletion on BRCA1 protein levels.

To this end, we used RNAi to deplete HIPK2 and analysed HIPK2 and BRCA1 protein levels in IR-damaged cells and in cells left untreated. Strikingly, immunoblotting revealed a clear drop in BRCA1 protein levels in the IR-treated HIPK2-depleted cells (**Fig.5a**). Unexpectedly, we also observed diminished BRCA1 protein levels in untreated HIPK2-depleted cells (**Fig.5a**), indicating that HIPK2 – in contrast to Pin1^36^ - does not exclusively regulate BRCA1 protein levels upon IR damage but also in undamaged cells. These findings suggest that HIPK2 can regulate BRCA1 through Ser1191 phosphorylation upon IR damage, and, in addition, through a Ser1191-independent mechanism in the absence of IR damage. Next, we used an additional HIPK2-specific siRNA to deplete endogenous HIPK2, which also resulted in a massive drop in BRCA1 protein levels already in undamaged U2OS cells (**Fig.5b**). Of note, protein levels of Abraxas, a BRCA1 complex protein^37^, were not reduced, excluding non-specific RNAi effects (**Fig.5b**). In addition, comparable results were obtained in HCT116 (**Fig.5c**) and RPE-1 cells (**Fig.5d**), where HIPK2 depletion also reduced BRCA1 protein levels and left Abraxas levels unaffected. These results exclude a cell line-restricted effect.

**Figure 5.**
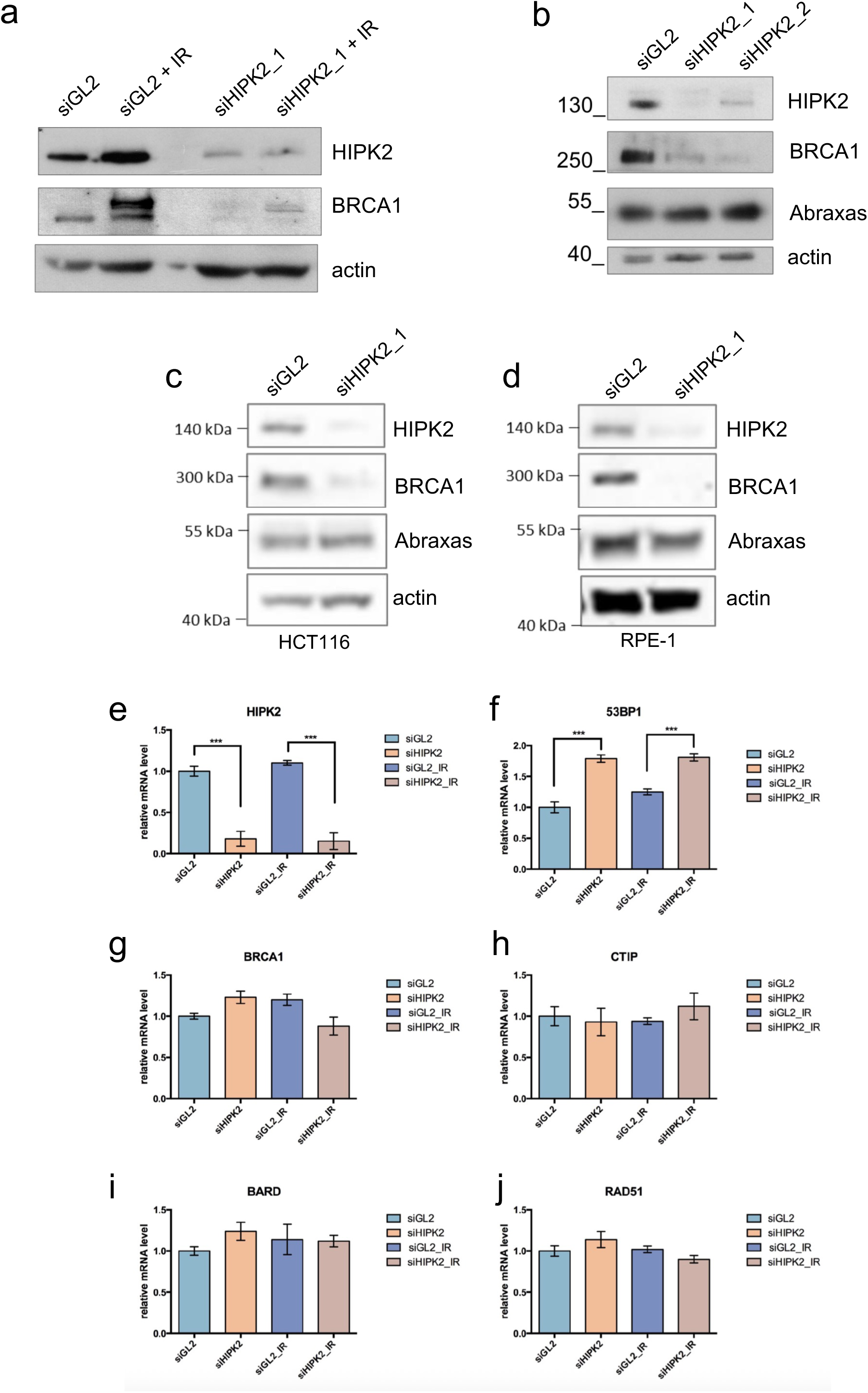
HIPK2 regulates BRCA1 protein levels. **(a)** HIPK2 depletion leads to a drop in BRCA1 protein levels in IR-damaged cells. Control (GL2) and HIPK2-specific RNAi was performed in U2OS cells. Subsequently, cells were treated with IR or left untreated as indicated, and 18 hours later immunoblot analysis was performed using the antibodies indicated. A representative experiment is shown (n=2). **(b)** HIPK2 depletion results in reduced BRCA1 levels in untreated cells. Control (GL2) and HIPK2-specific RNAi was performed in U2OS cells as indicated and 18 hours later immunoblot analysis was performed using the antibodies indicated. A representative experiment is shown (n=2). **(c,d)** HIPK2 depletion leads to reduced BRCA1 levels in untreated (c) HCT116 and in untreated (d) RPE-1 cells. Control (GL2) and HIPK2-specific RNAi was performed in U2OS cells as indicated and 18 hours later immunoblot analysis was performed using the antibodies indicated. A representative experiment is shown (n=2). **(e-j)** Reduced BRCA1 protein levels upon HIPK2 depletion do not originate from transcriptional effects. qRT-PCR analysis excludes transcriptional effects of HIPK2 depletion on BRCA1 and numerous other DSB repair factors. Representative experiments are shown (n=3 replicates).

In addition, qRT-PCR analysis was performed to investigate whether reduced BRCA1 levels may originate from a transcriptional effect exerted by HIPK2 depletion. While HIPK2 mRNA levels were, as expected, decreased upon HIPK2 RNAi (**Fig.5e**), no reduction in 53BP1 (**Fig.5f**), and also no reduction in BRCA1 mRNA levels (**Fig.5g**) was detected. Furthermore, no diminished mRNA levels of the resection factor CtIP (**Fig.5h**), BARD1 (**Fig.5i**), which is a stoichiometric BRCA1 binding partner, or Rad51 (**Fig.5j**) was observed. Together, these results further exclude unspecific transcriptional effects of the HIPK2 RNAi. In sum, these data indicate that HIPK2 regulates BRCA1 at post-transcriptional level and suggest a critical role for HIPK2 for regulating BRCA1 protein stability.

### HIPK2 depletion sensitizes cancer cells to IR

So far our results have demonstrated that depletion of HIPK2 diminishes BRCA1 protein levels in BRCA1-proficient cancer cells, and thus functionally mimics genetic loss or functional inactivation of BRCA1 (termed BRCAness), which is frequently observed in cancer including ovarian, breast and prostate cancer ^6^. Since HR-deficient cells have been reported to show increased radiosensitivity in cell culture ^38, 39^, we next analysed radiosensitivity of HIPK2-depleted cells by using colony formation assays. As expected, HIPK2 depleted (siHIPK2) cells showed an increased radiosensitivity when compared to the control (siGL2) cells (**Fig.6a,b**). Of note, already untreated HIPK2-depleted cells demonstrated reduced colony formation when compared to the control cells (**Fig.6c**), suggesting that the compromised HR repair results in reduced cell viability. In line with this interpretation, immunoblotting indicated that HIPK2-depletion leads to p53 stabilization already in the absence of IR damage (**Fig.6d**), presumably due to increased DNA damage because of the compromised HR. In sum, our results indicate increased sensitivity of HIPK2-depleted cancer cells to IR damage.

**Figure 6.**
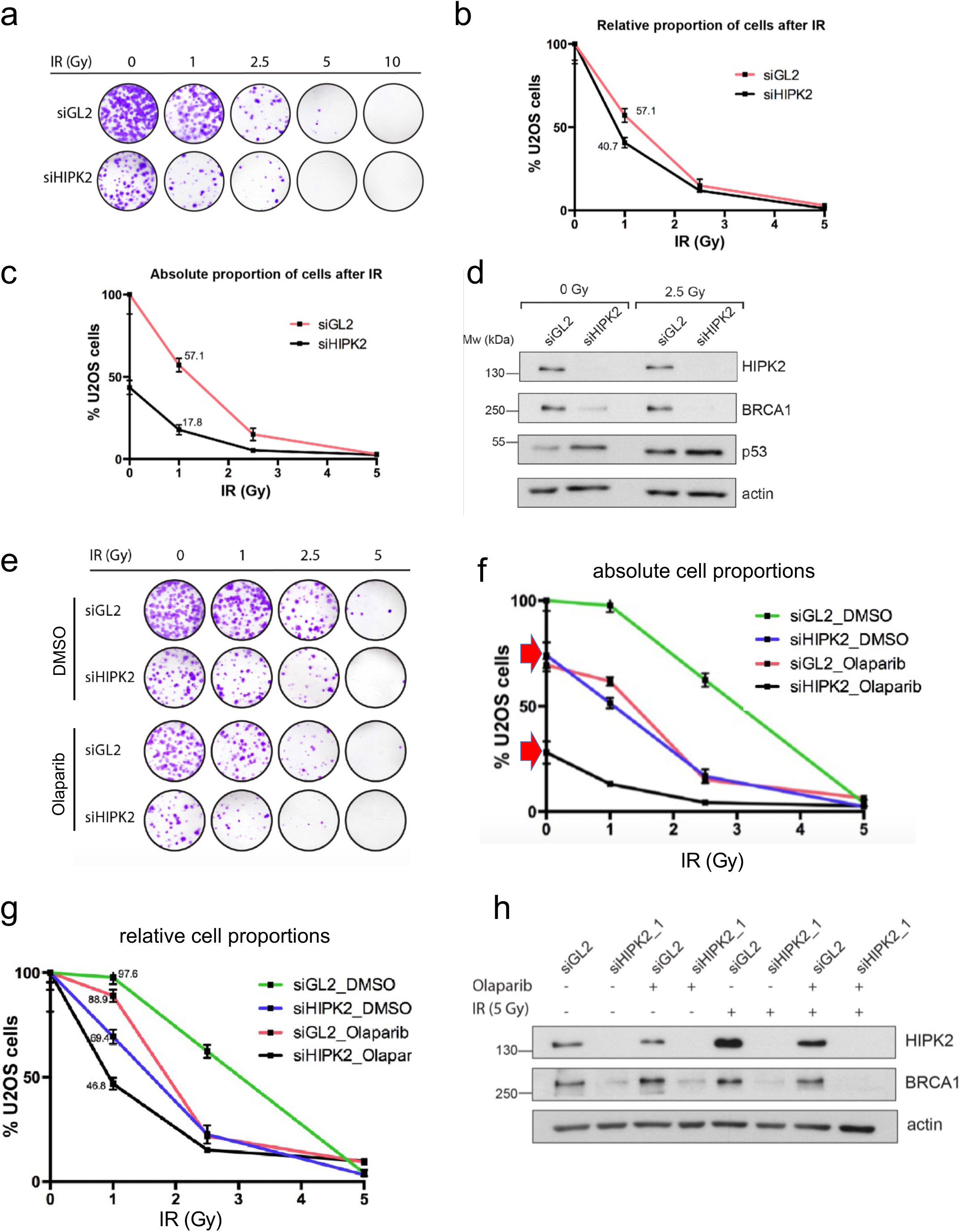
HIPK2 depletion sensitizes cells to IR damage, to PARP inhibition and combined PARPi and IR treatment. **(a)** HIPK2 depletion sensitizes to IR damage. U2OS cells were transfected with the indicated siGL2 and siHIPK2_1 as indicated. 24 hours later cells were treated with IR damage as indicated. Colony formation assays were fixed after 8 days and stained with crystal violet. Representative plates are shown. **(b)** HIPK2 depleted cells are hypersensitive to IR damage. Relative cell growth of cells after IR. Cell-bound dye from (a) was quantified at 570 nm, and values were normalized to the respective non-irradiated siGL or siHIPK2 samples. Mean and standard deviations are shown. (n=2, triplicates each). **(c)** HIPK2 depletion already reduces colony formation in untreated cells. Absolute cell proportion after IR damage. Cell-bound dye from (a) was quantified at 570 nm and normalized to the control (siGL2, untreated). Mean and standard deviations are shown. (n=2, triplicates each). (d) Immunoblotting verifies reduced BRCA1 levels upon HIPK2 depletion of the cells used for the colony formation analysis. U2OS cells transfected with siGL2 or siHIPK2_1 were irradiated with 2.5 Gy and analysed by immunoblotting as indicated. A representative experiment is shown (n=2). **(e, f, g)** HIPK2 depletion sensitizes for PARPi, and combined PARPi and IR treatment. U2OS cells were transfected with siGL2 and siHIPK2 as indicated. (e) Cells of the respective samples were seeded on 6-well plates, and four hours prior to irradiation media was supplied with either 100 µM Olaparib or the respective volume of the solvent DMSO. Cells were kept for 12-14 days and then stained with crystal violet (CV). Digital scans are shown. (f) Cell-bound CV was quantified and normalized to the DMSO/siGL2 treated sample (absolute cell growth). (g) Absorbance was normalized to the respective non-irradiated sample of each data series (relative cell growth). Mean values and respective SD are shown. (n=2, triplicates each). (f) HIPK2 knockdown efficiency was assessed by immunoblotting. Using the antibodies indicated. A representative experiment is shown (n=2).

### HIPK2 depletion sensitizes to PARPi and combined PARPi and IR treatment

BRCAness, or more general defective HR repair (HRDness), generates a cancer cell vulnerability that can be therapeutically exploited in the clinics due to its synthetic lethality with PARP inhibitor (PARPi) treatment ^6, 7, 40^. To this end, we analysed whether HIPK2-depletion, which results in defective HR, sensitizes BRCA1-proficient cancer cells to treatment with the PARPi Olaparib, and whether HIPK2-depleted cancer cells shown increased sensitivity for combined PARPi and IR treatment. Colony formation assays displaying absolute colony numbers revealed an increase sensitivity of HIPK2-depleted cells to treatment with PARPi Olaparib (**Fig.6e,f**, compare red arrow heads in f). In addition, we also observed that PARPi treatment robustly increased radiosensitivity of HIPK2-depleted cells (**Fig.6e,g**). Immunoblotting confirmed reduced BRCA1 protein levels upon HIPK2 depletion in the treated cancer cell (**Fig.6h**). Our results indicate that HIPK2 depletion leads to increased PARPi sensitivity and mild PARPi treatment increases cancer cell radiosensitivity.

### Pharmacological HIPK2 inhibition triggers PARPi hypersensitivity of BRCA1-proficient cancer cells

Since HIPK2 is Ser/Thr kinase, we also aimed at using pharmacological HIPK2 inhibition to investigate potential effects on PARPi sensitivity in BRCA1-proficient cancer cells. A recent publication demonstrated strong and specific inhibitory effects of Abemaciclib on the activity of HIPK protein kinases family members including HIPK2^8, 41^. To test the effect on HIPK2 inhibition in cells, we first tested whether treatment with Abemaciclib impacts on HIPK2 and BRCA1 protein levels using immunoblotting. Actually, Abemaciclib treatment resulted in reduced steady-state HIPK2 protein levels in the absence and in presence of 2 Gy IR (**Fig.7a**), which is expected and in line with our previous findings that HIPK2 activity is critical for its protein stability^21^. Notably, Abemaciclib treatment also resulted in reduced BRCA1 levels (**Fig.7a**), similar to that we observed upon HIPK2 depletion. These results motivated us to analyse the effect of pharmacological HPK2 inhibition using Abemaciclib on cellular PARPi Olaparib sensitivity (**Fig.7b**). Strikingly, HIPK2 inhibition with Abemaciclib significantly increased PARPi sensitivity of BRCA1-proficient U2OS cancer cells (**Fig.7c,d**). Comparable results were obtained in BRCA1-proficient HCT116 colon cancer cells (**Fig.7e,f**). In sum, these results shown that pharmacological HIPK2 inhibition established a BRCAness phenotype in BRCA1-proficient cells, and thus is synthetic lethal with PARPi treatment.

**Figure 7.**
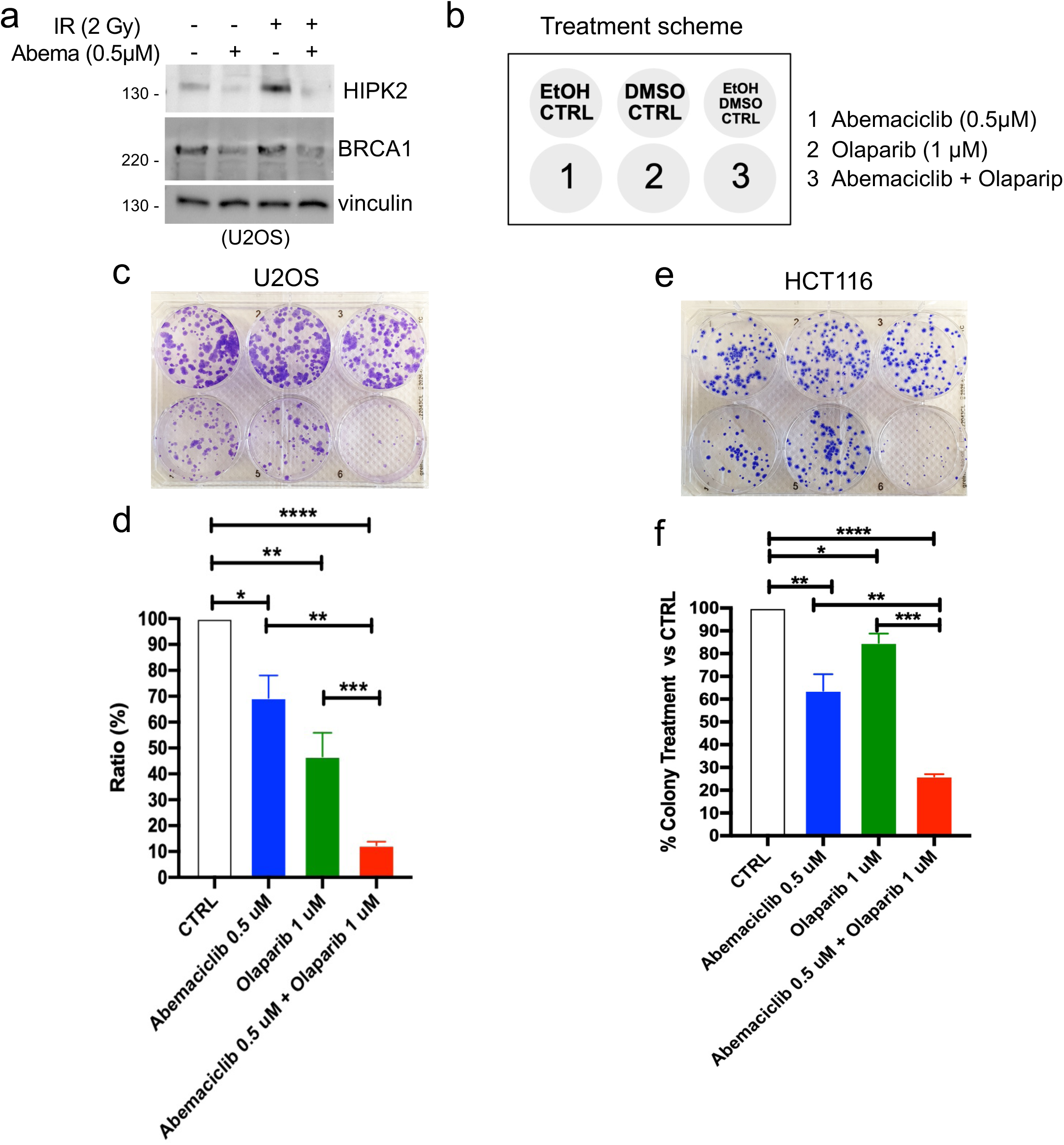
Pharmacological HIPK2 inhibition triggers PARPi hypersensitivity in BRCA1-proficient cancer cells. **(a)** Pharmacological inhibition of HIPK2 diminished BRCA1 levels in BRCA1-proficient cancer cells. U2OS cells were treated with the indicated doses of Abemaciclib and Olaparib (PARPi). After 18 hours cells were analysed by immunoblotting as indicated. A representative experiment is shown (n=3 biological replicates). **(b)** Treatment scheme for the analysis performed in (c,d) U2OS and (e,f) HCT116 cells. **(c, d)** Pharmacological HIPK2 inhibition sensitizes U2OS cells to PARPi treatment. Cells were treated with Abemaciclib and Olaparib as indicated in (b), and colony formation assays were performed. **(c)** A representative colony formation assay data set is shown. **(d)** Quantification of the colony formation data of n=3 biological replicates. **(e, f)** Pharmacological HIPK2 inhibition sensitizes HCT116 cells to PARPi treatment. Cells were treated with Abemaciclib and Olaparib as indicated in (b), and colony formation assays were performed. **(e)** A representative colony formation assay data set is shown. **(f)** Quantification of the colony formation data of n=3 biological replicates.

## DISCUSSION

The Ser/Thr protein kinase HIPK2 is a well-established regulator of the cell death-inducing p53 response in the wake of irreparable DNA damage. However, whether HIPK2 plays a role in response to repairable DNA damage, and in particular in DSB repair, has remained elusive. Here, we discovered an unforeseen role of HIPK2 in the regulation of HR and PARPi sensitivity.

We found that HIPK2 is stabilized and activated in response to repairable DSB damage to a similar extend as previously reported in response to irreparable damage^16, 18, 19, 21, 26, 42^, which suggested a currently undefined function in the regulation of DSB repair. Accordingly, no molecular markers of DNA damage-induced apoptosis were observed in cells demonstrating HIPK2 activation upon repairable IR damage. Moreover, we found that HIPK2 accumulates at DSBs upon IR damage. HIPK2 was detected at DSBs using both a pan-specific HIPK2 antibody, and a HIPK2 antibody targeting the autophosphorylated, active pT880/pS882-phosphorylated HIPK2^21^. Active HIPK2 partially colocalized with a set of DNA damage foci markers including γH2AX, Ser1981-phosphorylated ATM, 53BP1 and BRCA1. Pharmacological inhibition of ATM prevented accumulation of HIPK2 at DSBs, which is in line with our previous findings demonstrating a critical role of ATM in HIPK2 activation and stabilization upon induction of DSBs^16, 18, 21^.

Evidence that HIPK2 regulates HR activity came from reporter-based repair assays assessing DSB repair through HR and the NHEJ pathway, which revealed a defect in HR repair upon HIPK2 depletion. NHEJ was not impaired. Furthermore, analysis of sister-chromatid exchanges (SCEs), which are formed by homologous recombination activity, illustrated a significant reduction in SCE numbers in cells depleted for HIPK2, pointing to defective HR activity in HIPK2-depleted cells. Furthermore, a robust reduction in IR-induced Rad51 foci formation and in BRCA1 foci formation was detected upon HIPK2 knock-down, which pointed to a potential interplay between BRCA1 and HIPK2.

Interaction analyses indicated BRCA1-HIPK2 complex formation in cells, and *in vitro* analysis using GST pulldown assays revealed physical interaction of BRCA1 and HIPK2, further strengthening our hypothesis that HIPK2 can impact on the HR pathway through regulating BRCA1. Along these lines, HIPK2 kinase assays identified Ser1191 as a major BRCA1 phosphorylation site, and generation of a phospho-specific BRCA1 pSer1191 antibody enabled us to demonstrate accumulation of pSer1191 BRCA1 upon IR damage at DSBs. Ser1191 has been previously reported to play a critical role for BRCA1 stability after IR damage by a mechanism involving BRCA1 isomerization through the phospho-specific cis/trans isomerase Pin1, which binds to pSer1191 ^36^. In light of this report, and supported by our qRT-PCR analysis demonstrating no altered BRCA1, BARD1, CtIP and Rad51 mRNA expression levels upon HIPK2 depletion, we propose that HIPK2 regulates BRCA1 at post-transcriptional levels through regulation of its protein stability. Interestingly, our previous work identified Pin1 as a critical regulator of DSB-induced HIPK2 stabilization through binding HIPK2 autophosphorylated at Thr880/Ser882.^21^, placing Pin1 at center stage in IR-induced BRCA1 regulation through controlling stability of the IR-activated BRCA1 Ser1191 kinase and of BRCA1 itself.

In contrast to Pin1 which mediates BRCA1 stability exclusively in response to IR damage^36^, our results indicate that depletion of the IR-induced BRCA1 Ser1191 kinase HIPK2 not exclusively affects BRCA1 levels in response to IR damage, but also results in reduced BRCA1 steady-state protein levels in the absence of IR damage. These results suggest that HIPK2 regulates BRCA1 protein levels - beyond BRCA1 Ser1191 phosphorylation upon IR damage - through an additional mechanism which remains to be uncovered in the future. Whether here additional HIPK2 phosphorylation events are involved, such as the *in vitro* phosphorylation sites mapped *in vitro* to the very C-terminal stretch of BRCA1 comprising aa1491 to 1863 (**Fig.3a**) will be clarified in the future. In addition, whether the steady-state BRCA1-HIPK2 complex found in undamaged cells also contribute to HR repair, or might be linked to additional BRCA1 functions such as balancing transcription-coupled stress^43^, remains to be elucidated in the future.

Our results indicate that HIPK2 depletion or its pharmacological inhibition results in reduced BRCA1 levels in BRCA1-proficient cells, mimicking BRCAness, which is a frequently observed HR defect in cancer cells that can be therapeutically exploited due to synthetic lethal effects with PARP inhibitors^7^. Accordingly, HIPK2 depletion and pharmacological HIPK2 inhibition using Abemaciclib ^41^, resulted in increased PARPi sensitivity of BRCA1-proficient cancer cells. Although Abemaciclib is a *bona fide* Cdk4/6 inhibitor, recent data suggest an even higher affinity of this inhibitor for HIPK and DYRK family kinases^41^. However, although Abemaciclib inhibits IR-induced HIPK2 stabilization and reduces BRCA1 protein levels, we cannot exclude that these effects are mediated -at least in part-through its CDK4/6 inhibitory activity. Therefore, it will be interesting to develop (more) specific HIPK2 inhibitors in the future.

In summary, we propose the following scenario of how HIPK2 regulates the HR pathway in response to repairable IR damage (**Fig.8**): HIPK2 and BRCA1 form a complex in undamaged cells. In response to repairable IR damage, ATM activity mediates the stabilization and activation of HIPK2^16, 18^ and targets it to DNA damage foci formed at DSBs. Activated HIPK2 phosphorylates BRCA1 at Ser1191 and thereby mediating IR-induced, Pin1-mediated BRCA1 stabilization^36^, thus facilitating BRCA1 function in the homology-directed DSB repair pathway through CtIP-mediated DNA-end resection and Palb2-BRCA2 mediated Rad51 loading. Depletion of HIPK2 leads to HR-deficiency through destabilization of BRCA1, thus sensitizing cancer cells to PARP inhibitor treatment. In summary, our results suggest HIPK2 inhibition as a novel strategy to sensitize BRCA1-proficient cancer cells to PARPi treatment.

**Figure 8.**
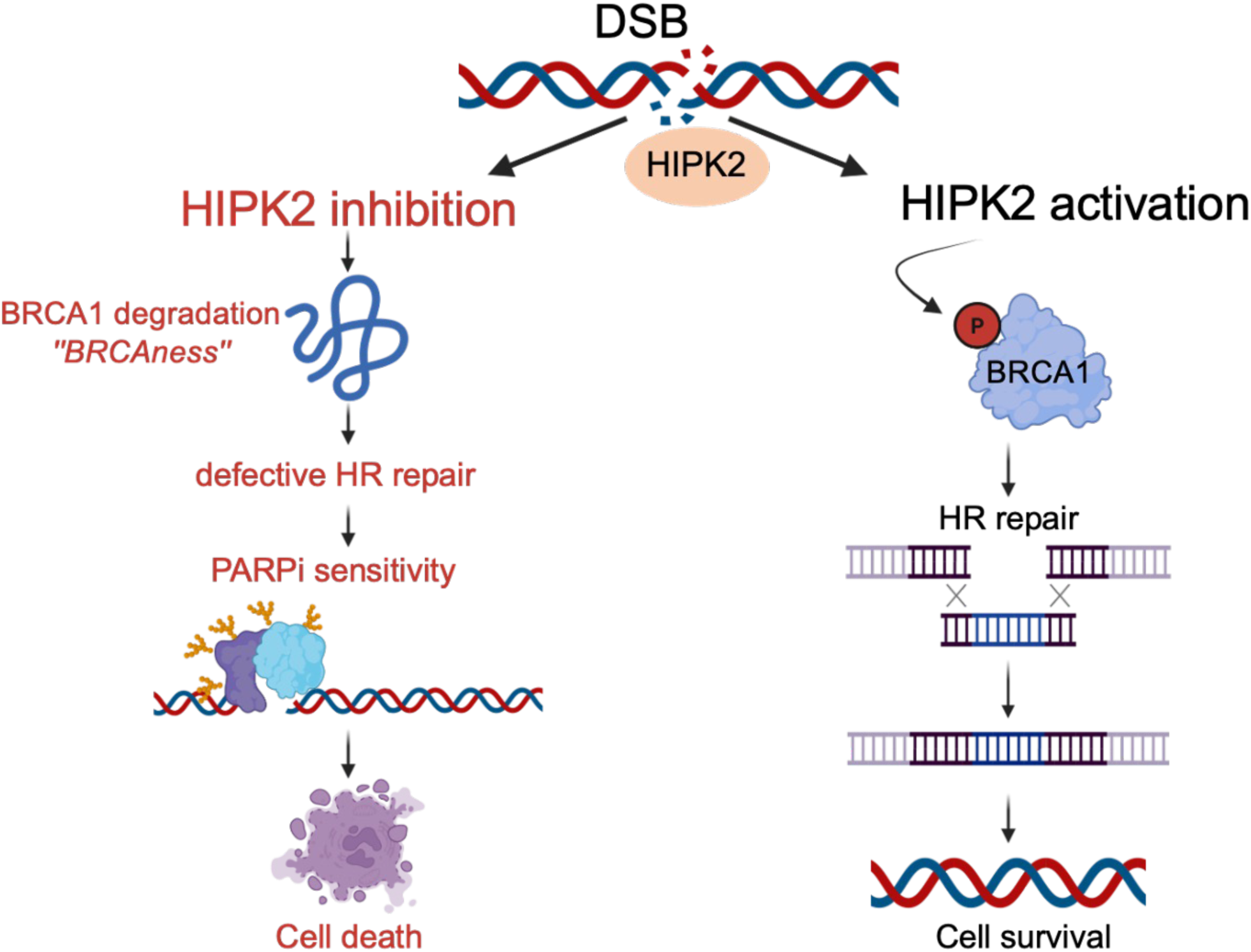
Proposed model for the role of HIPK2 in homology-directed DNA repair and PARPi sensitivity. Upon DSB formation HIPK2 is stabilized and activated in an ATM-dependent manner and accumulates at DSB. HIPK2 interacts with BRCA1 and phosphorylates BRCA1 at Ser1191, mediating BRCA1 stability and facilitating productive HR repair. Upon HIPK2 depletion or pharmacological inhibition, BRCA1 protein levels massively drop and leads to a BRCAness phenotype in BRCA1-proficient cancer cells, thus generating hypersensitivity for PARPi treatment. This graphics has been generated using BioRender.

## METHODS

### Cell lines, cell culture, transfection and cell treatment

U2OS, HCT116, RPE-1 (all obtained from ATCC) and RPE-1 BRCA1 +/+ and RPE-1 BRCA1 -/- (both in p53 KO background, kindly provided by Daniel Durocher^35^) were maintained in DMEM (Gibco) supplemented with 10% FCS, 1% (w/v) penicillin/streptomycin, 2 mM L-Glutamine, 1mM Sodium Pyruvat and 20 mM Hepes buffer at 37°C at 5% CO_2_. Transient transfections were done using Lipofectamine 2000 (Invitrogen) or by standard calcium phosphate precipitation. For cell treatment, cells were incubated with culture medium supplemented with Doxorubicin, Olaparib, Abemaciclib at the specified concentrations and subsequently harvested.

### BRCA1 pSer1191 antibodies

The phospho-specific BRCA1 Ser1191 antibodies were generated by immunizing rabbits with the following KLH-coupled peptide: NH_2_-CLSRSP(pS)PFTH-CONH_2_. Rabbit sera were affinity-purified against the phospho-peptide and subsequently non-phosphorylation specific antibodies were removed by a column containing the immobilized corresponding non-phospho-peptide. Subsequently, antibody specificity has been validated for immunoblotting and immunofluorescence staining.

### Sister-chromatid exchange (SCE) assay

SCEs have been performed essentially as previously published^44, 45^. In brief, cells transfected with the siRNA indicated were grown for the duration of one cell cycle in MMS in the presence of the thymidine analog BrdU (Sigma-Aldrich). BrdU was added at a final concentration of 10 µg/ml to fresh culture medium immediately after the pulse treatment with the methylating agent. The cells were incubated strictly in the dark. Chromosome preparations were performed, air-dried and after 1–2 days differentially stained according to a modification of the original FPG-technique^46^. At least twenty-five metaphases were evaluated per treatment level. Only fully differentiated metaphases were evaluated.

### DR-GFP (HR) and pimEJ5GFP (NHEJ) reporter assays

For determining the influence of HIPK2 depletion on homology-directed repair the DR-GFP^32^ (Addgene plasmid #26475) assay was used, and for alternative NHEJ the pimEJ5GFP^31^ (Addgene plasmid #44026) was used. U2OS cells were generated that contain stabile genomic integrated DR-GFP and pimEJGFP reporter constructs, respectively, and the assays were performed as previously described. ^31, 32^ In brief, U2OS DR-GFP or U2OS-pimEJ5GFP cells was knocked down using RNAi, and the influence on homologous recombination (DR-GFP cells) or alternative NHEJ (pimEJ5GFP cells) was determined by transfection with a plasmid that expresses the rare cutting endonuclease I-SceI^47^ (pCBASceI, Addgene plasmid #26477) as described. GFP signals, showing homologous recombination repair activity for U2OS DR-GFP and alternative end-joining for U2OS-pimEJGFP cells, were quantified by flow cytometry (FacsCantoII, BD Bioscience).

### Expression constructs

All human HIPK2 constructs have been described previously^21, 23^ ^16^. Human HA-BRCA1 cDNA was obtained from Addgene (plasmid #14999), and all used BRCA1 construct were generated by standard PCR techniques and molecular cloning. All constructs were verified by DNA sequencing.

### GST pulldown assays

GST fusion protein expression, protein purification and GST-pulldown assays were performed essentially as previously published^16, 21^. In brief, GST fusion proteins were expressed in *E.coli* BL21 and purified using glutathione (GSH) sepharose 4B beads (GE Healthcare). 2 µg of bead-coupled GST fusion proteins were used for the pulldowns. GST fusion proteins were incubated with indicated human HIPK2 protein generated by *in vitro* translation using the TNT Coupled Reticulocyte Lysate System (Promega) and ^35^S-labelled methionine according to the manufactureŕs instructions. GST-pulldowns were performed in 1x PBS, 0.05% NP40, 1 mM PMSF, 1 mM Na_3_VO_4_ and analyzed by SDS-PAGE, Coomassie Brilliant Blue staining and autoradiography of the dried gels.

### Immunoblotting and immunoprecipitation

Immunoprecipitation analysis and immunoblotting was performed in principle as previously published^16, 21^, and proteins were detected by enhanced chemiluminescence (SuperSignal West Dura and Femto, Pierce). For coimmunoprecipitation cells were lysed in lysis buffer (20 mM Tris–HCl pH 7.4, 10% glycerol, 1% NP40, 150-250 mM NaCl, 25 mM NaF, 5 mM EDTA, 1 mM sodium vanadate). Immunoprecipitation (IP) of endogenous HIPK2 was performed as previously described by us^16^. IP reactions were incubated for 1-4 h at 4°C on a rotating wheel and washed three times in lysis buffer. Immunocomplexes were boiled at 95°C for 5 min, separated by SDS-PAGE and analysed with specific antibodies. The following primary antibodies were used for immunoblotting or IP: p53 (DO-1) from Santa Cruz Biotechnologies, actin (C4) from MP Biomedicals, BRCA1 (D-9, Santa Cruz, sc-6954), PARP (4C10-5) from BD Pharmingen, p53 phospho-Ser46 and cleaved PARP from Cell Signaling Technologies, and AKT (Mab, 9Q7) from Invitrogen. The affinity-purified rabbit pan-specific HIPK2 antibodies^16, 23^ and the phosphorylation-specific pThr880/pSer882 HIPK2 antibody^21^ have been described previously.

### Immunofluorescence staining and confocal microscopy

Indirect immunofluorescences were performed in principle as published. In brief, cells were fixed with 4 % PFA for 10 minutes, permeabilized with 0,5 % Triton and stained with indicated antibodies. As secondary antibodies Alexa Fluor^®^488 or 594 (Life technologies) goat anti-rabbit and goat-anti mouse, or Cy3 goat anti-mouse antibody (Dianova) were used. DNA was visualized with Hoechst (Sigma Aldrich) or TO-PRO-3 (Thermo Fisher Scientific) staining. The following primary antibodies used for immunofluorescence staining (IF): pATM (Ser1981) (10H11.E12, Cell Signalling) dilution 1:100 ; Rad51 (Abcam, ab63801) dilution 1:2000; phospho-histone H2AX Ser139 (Mab, 20E3, Cell Signaling) dilution 1:250; BRCA1 (D-9, Santa Cruz, sc-6954) dilution: 1:50; BRCA1 pSer1191 (self-generated, rb polyclonal) dilution 1:200; 53BP1 (clone BP13; Sigma Aldrich, Mab3802) dilution 1:250, pT880/S882 HIPK2^21^ (self-generated rb polyconal) dilution 1:100. Confocal microscopy was performed by using a LSM 710 system (Zeiss) or using the Olympus FluoView FV1000 IX-81 (Olympus) confocal microscope. Image analysis and quantification were performed using Fiji and GraphPad Prism, and for FV1000 IX-81 by using the corresponding software FV10-ASW 2.0 and ImageJ.

### RNA interference

siRNA duplexes were produced by Sigma Aldrich and Ambion. For HIPK2 knockdown the following siRNAs were used: siHIPK2_1 (Sigma Aldrich): 5’-CCAGGUGAACAUGACGACAGA-3’, siHIPK2_2 (Sigma Aldrich): 5’-GCGUCGGGUGAAUAUGUAU-3’ , siGL2 (Sigma Aldrich) : 5’-AACGUACGCGGAAUACUUCGAdTdT-3’, BRCA1 (Ambion Silencer Select): 5’-CAUGCAACAUAACCUGAUAdTdT-3’, siCTRL (AmbionSilencerSelect, Art.No.: 4390844). siRNA duplexes (final concentration 20-50 nM) were transfected using Lipofectamine RNAiMAX (Life Technologies) or HiPerFect (Qiagen) as specified by the manufacturer.

### Flow cytometry

To determine cell cycle distribution and sub-G1 DNA content, cells were stained with PI (30 µg/ml in PBS) and cell cycle distribution was determined by flow cytometry. using a BD FACSCanto™ II. Experiments were repeated at least three times and a representative example is shown.

### Quantitative real-time PCR (qRT-PCR)

RNA was isolated using the TRIzol® reagent (Invitrogen) according to the manufacturer’s instructions and 0,5 to 1 µg RNA was reverse transcribed according to manufacturers’ protocols (Applied Biosystems, High Capacity RNA to cDNA Kit). To avoid amplification of genomic DNA RNA was treated with RNase free DNase (Roche). Quantitative real-time PCR (Real Time PCR System 7300, Applied Biosystems) was carried out using pre-designed FAM and MGB labelled TaqMan^®^ primer (GAPDH 4333768-1005027, HPRT1 4333764-0912033, BRCA1 Hs01556193_m1, BARD1 Hs00184427_m1, HIPK2 Hs00179759-m1, CtIP Hs00161222_m1, 53BP1 Hs00996818_m1, Rad51 Hs00947967_m1) and TaqMan^®^ Gene Expression Master Mix according to the manufacturers’ protocol (Applied Biosystems). Results were analysed using the ΔΔCt-method. Here, samples were normalised against the housekeeping genes GAPDH or HPRT1, and finally illustrated using the GraphPad Prism software.

### Colony formation assays

Colony formation assays were used to analyse clonogenic cell growth after treatments. For the experiments on the effect of pharmacological HIPK2 inhibition we used the following protocol: 500 and 1000 U2OS or HCT116 cells were seeded in 6-well plates and treated at the same time with Abemaciclib 0.5 µM (Sigma-Aldrich) and Olaparib 1 µM (Selleckchem). After 96h the medium was changed. After 10 to 14 days in culture, The cells were fixed with methanol 100% for 20 min at room temperature and colored with crystal violet solution (0.2% (w/v) crystal violet, 5% (v/v) acetic acid, 5% ethanol (v/v)) for 5 min at room temperature. The excess of staining solution was washed using distilled water. Colonies were counted manually. For the experiments shown in Figure 6, 1000 U2OS cells were seeded in 6-well plates in DMEM complete. Plates were kept in cell culture for 12-14 days. Growth media was changed once after 8 days. For crystal violet (CV) staining, media was removed, cells washed twice with PBS and 1 ml staining solution (0.2% (w/v) crystal violet, 5% (v/v) acetic acid, 5% ethanol (v/v)) added. After 10 min incubation at room temperature, the staining solution was removed and plates washed three times with tab water. Plates were dried upside down overnight and scanned to generate digital images. To quantify the amount of cell-bound dye, crystal violet was resuspended in 1 % SDS solution by shaking at room temperature for at least 8 hours. The absorption at 570 nm was measured using a photometer (plate reader) and analyzed

### *In vitro* HIPK2 kinase assays

*In vitro* HIPK2 kinase assays for substrate phosphorylation have been performed as previously published^19, 21^. In brief, 0.2 - 0.5 µg bacterially expressed and purified 6xHis-HIPK2 protein was incubated in 30 µl kinase buffer containing 40 µM cold ATP and 5 µCi [γ-^32^P] ATP and 1-2 µg of the indicated GST-BRCA1 substrate proteins as described. Incubation with Myelin Basic Protein (MBP, Sigma) was used as a control substrate as described previously^19, 21^. After incubation for 30 min at 30°C, the reaction was stopped by adding 5x SDS loading buffer. After separation by SDS-PAGE, gels were fixed, dried and exposed to X-ray films.

### Statistical analysis

All experiments in this study were independently repeated three times (biological replicates). Quantitative data are presented as the mean ± standard deviation. Statistical analyses were performed to evaluate the significance of differences between experimental groups. Since datasets consisted of matched sample pairs measured under different experimental conditions, statistical significance was assessed using a two-tailed paired Student’s T-test. This test was chosen to account for the inherent pairing of samples, minimizing inter-experimental variability and allowing accurate assessment of condition-specific effects. A *p*-value of less than 0.05 was considered statistically significant. Significance levels are represented as follows: *, p < 0.05; **, p < 0.01; ***, p < 0. 001. All statistical analyses and graphical visualizations were carried out using GraphPad Prism software.

## Acknowledgements

We are grateful to Dan Durocher for the kind gift of BRCA1 +/+ and BRCA1 -/- RPE-1 cells. This work has been supported by the Deutsche Forschungsgemeinschaft, SFB 1361 (Project-ID 393547839), project 19 (to TGH).

## Declaration of interests

The authors declare no competing interests.

## Author contributions

P.W., P.-O.F., G.N., S.K., W.P.R, T.N., K.R., S.K., M.C.L., S.D., D.P., M.C., M.A., and T.G.H. designed the experiments. P.W., P.-O.F., G.N., S.K., W.P.R., T.N., D.P., S.K., H.B. and Y.H. performed the experiments. S.K., T.N., S.D., W.P.R., M.C.L., G.N., M.A., and P.-O.F., analyzed the data. T.G.H. wrote the manuscript, and all authors commented on the manuscript. T.G.H.. supervised the research.

## Notes

### Competing Interest Statement

The authors have declared no competing interest.

### Summary of Updates

In this revised version several typos identified in the manuscript and in the figures were corrected, and a missing antibody reference in the Materials section has been added.

